# Development and Validation of the *Vanderbilt PRS-KS*, an Instrument to Quantify Polygenic Risk Score Knowledge

**DOI:** 10.1101/2022.10.26.513776

**Authors:** Doug Stubbs, Gillian Hooker, Yajing Le, Lucas Richter, Alexander Bick

**Author notes:** **Corresponding Author**, Doug Stubbs, MGC, |.

## Abstract

**Purpose:** As polygenic risk scores (PRS) enter clinical practice, healthcare providers’ and the publics’ comprehension of PRS results are of great importance, yet poorly understood. We present the Vanderbilt Polygenic Risk Scores Knowledge Score (Vanderbilt PRS-KS), a tool to quantify PRS knowledge.

**Methods:** The Vanderbilt PRS-KS was developed by a team of genetic counselors and physicians to cover key conceptual facts pertaining to PRSs. We recruited (n=500) individuals with demographics representative of a U.S. sample and graduate-level healthcare students (n=74) at a large academic medical center to participate in this validation study. We evaluated the Vanderbilt PRS-KS’s psychometric properties using confirmatory factor analysis (CFA) and item response theory (IRT).

**Results:** The 7-item Vanderbilt PRS-KS correlated to a single latent construct on CFA (Λ=0.31-0.61). The scale showed promising reliability (Cronbach’s α=0.66) with IRT summed scores of ≥2 to ≤5, demonstrating reliability > 0.70. The Vanderbilt PRS-KS significantly correlated with genetic knowledge and applied PRS knowledge (r=0.55, r=0.29), and graduate-level healthcare students had significantly higher scores compared to the representative sample (p<0.01).

**Conclusions:** The Vanderbilt PRS-KS is a rigorously validated measure to quantify PRS knowledge.

## Introduction

Complex disorders are caused by alterations in many genes with small effect sizes, often coinciding with environmental factors, that influence an individual’s risk of developing disease over time.^1^ Genome-wide association studies elucidate variant associations in many genes to outcomes of interest, which researchers quantify and aggregate to form polygenic risk scores (PRS).^2^ A polygenic risk score is a sum of the established risk alleles, weighted by effect size, on an outcome. PRSs can stratify an individual’s disease risk in combination with traditional risk factors (i.e., family history, age, smoking, etc.) or as a potential standalone tool when monogenic and traditional risk factors are insufficient or poorly understood.

As PRSs enter clinical practice, healthcare providers’ and patients’ comprehension of results and the proper risk communication strategies are of great importance yet poorly understood. A validated PRS knowledge measure could identify current gaps in knowledge, elucidate potential risks and benefits, promote responsible PRS use, and serve as an objective tool for future research. We present the Vanderbilt Polygenic Risk Scores Knowledge Scale (Vanderbilt PRS-KS), a tool to quantify PRS knowledge. Our goal was to develop and validate a measure with the following criteria: 1) encompass benefits and limitations of PRSs, 2) utilize context-neutral language, and 3) demonstrate sound psychometric properties. To our knowledge, no existing PRS knowledge measure fulfills these criteria.

## Materials and Methods

### Respondents and Procedures

The Vanderbilt PRS-KS was validated in 500 individuals recruited through prolific.co, an online research and survey distribution platform.^3^ The 500 individuals were representative of U.S. Census Bureau data for age, sex, and ethnicity. Additionally, 75 graduate students in the Vanderbilt University School of Medicine (VUSM) and Vanderbilt University School of Nursing (VUSN) were recruited to assess for differences in Vanderbilt PRS-KS functioning compared to the U.S. representative sample. VUSN and VUSM students were ascertained through recruitment fliers and word of mouth. Study data were collected and managed using REDCap electronic data capture tools hosted at Vanderbilt University.^4,5^ The institutional review board of the Vanderbilt University Medical Center approved the study’s protocol

### Survey Measures

#### The Applied PRS Knowledge Measure

The Applied PRS Knowledge Measure was derived from commercial lab reports. The 3-item Applied PRS Knowledge Measure assesses the individual’s ability to interpret a graphical PRS result and the individual’s understanding of the limitations of the results.

#### Demographics

Demographics were self-reported and included sex, race, and ethnicity.

#### Genetic Knowledge

The previously validated 19-item UNC_GKS was used to assess individuals’ genetic knowledge.^6^ The UNC_GKS genetic knowledge includes statements covering genes, genes and health, and how genes are inherited.

#### Attitudes Towards Genetics

Attitudes towards genetics were measured using the 6-item ‘favorable attitude’ subscale of Morren et al.’s Genetic Attitudes Scale.^7^

### Data Analysis

Descriptive statistics and psychometrics of the Vanderbilt PRS-KS were examined. The scale’s psychometric analysis included internal consistency-quantifies the relationship between a set of items, confirmatory factor analysis (CFA)-confirms a single underlying construct in the scale, item response theory (IRT)-investigates the relationships between individual responses to each item and the individual’s ability (PRS knowledge), and convergent validity-evaluates the expected associations with conceptually related variables.

### Item-Level Descriptive Statistics

We found the proportion of individuals correctly answering each item of the Vanderbilt PRS-KS to determine if there were floor effects (< 90% answering an item correctly) or ceiling effects (> 90% answering an item correctly), indicating items are too hard or too easy. Inter-item tetrachoric and item-total correlations verified positive correlations between items before beginning CFA.^8^ Receiving a negative or low item-total or inter-item correlation or violating floor or ceiling effects could indicate items are not measuring the same latent construct, and items may need to be removed or altered.

### Confirmatory Factor Analysis

Using R package lavaan,^9^ one-factor CFA was performed using the maximum likelihood (ML) estimator.^10^ Following suggested reporting guidelines,^11^ acceptable model fit was defined as chi-squared with a p-value > 0.05, Tucker-Lewis index (TLI) and comparative fit index (CFI) > 0.95, root mean square error of approximation (RMSEA) < 0.05, standardized root mean square residual (SRMR) < 0.8, and residual values modification indices < 10. Modification indices were used detect areas of local dependence, and items displaying local were considered for removal.^12^

Standardized factor loadings provide evidence of good item fit to a single underlying construct. Acceptable standardized factor loadings were defined as Λ > 0.30, items with loadings Λ < 0.30 were removed, and CFA was repeated with the remaining items.^13^

### Item Response Theory

There was three possible IRT model fits for the Vanderbilt PRS-KS, the one-, two- or three-parameter logistic IRT model (1PL, 2PL, or 3PL, respectively). The −2*log-likelihood of the two model calibrations, e.g., 1PL vs. 2PL, were used to determine model fit with the null hypothesis that the more parsimonious model fits well.^14^ A significant p-value indicates the more parsimonious model does not fit the data and more parameters would better fit the data. Item fit was assessed using the Orlando and Thissen’s S-χ statistic with a (P > 0.05) representing good fit.^15^

IRT has four key assumptions-scale unidimensionality, locally independent items, monotonicity, and item invariance.^14^ CFA confirmed the scale’s unidimensionality. Monotonicity (the assumption that as ability increases, the probability of a correct response increases) was confirmed by the s-shaped curves on the item characteristic curves (ICC). Modification indices < 10 provided evidence for local dependencies. Item invariance was examined through differential item functioning (DIF).^16^

DIF was examined using age, sex, race, and ethnicity variables in the 500 U.S. representative sample. Logistic regression models were used to look for a relationship between item responses and subgroups. The first model contained one predictor, IRT scores. The second model contained the IRT score predictor and a group membership predictor. The third model included the IRT score predictor, group membership predictor, and included the interaction between them. DIF was evaluated using a likelihood ratio test comparing model 1 and model 2 and nonuniform DIF by comparing models 2 and 3. After multiple comparisons, the Benjamini-Hochberg (BH) procedure with a false discovery rate (FDR) of 0.05 was applied to determine the presence of significant DIF.^17^

Similar to previous scale development methods, items with CFA or IRT misfit, violated local dependence, or exhibited DIF were removed.^6^ The remaining items were used to calculate the IRT test information function (TIF) and reliability.

### Item Response Theory Scoring and Reliability

The IRT test information function (TIF) curve was generated using the sum item’s information. Higher information represents higher precision at different levels of PRS knowledge.^18^ Typically, IRT scores have a mean of 0 with a standard deviation of 1. To enhance interpretability, summed IRT scores were scaled to the T-score metric, making the mean 50 with a standard deviation of 10.^6^ Each item response was weighted by its discrimination parameter (*a*), and items with greater discrimination have a more significant impact on the IRT scores. However, multiple combinations of correct and incorrect item responses scores can generate a summed score > 0, regardless of the scaled scores. A scoring table was developed, converting summed scores to expected scale T-score metrics.

### Classical Test Theory Reliability

Internal consistency of the Vanderbilt PRS-KS was computed using Cronbach’s α and the greatest lower bounds (GLB).^19,20^ Traditionally, acceptable Cronbach’s α values are ≥ 0.70, and poor values are < 0.60.^21^

### Convergent Validity

Pearson correlations were calculated between the Vanderbilt PRS-KS and the UNC-GKS (genetic knowledge), the Applied PRS Knowledge Measure, and the individuals’ Attitudes Towards Genetics. Positive, significant correlations between individuals’ genetic knowledge, applied knowledge, and attitudes towards genetics and Vanderbilt PRS-KS scores were anticipated.

Multivariate regression was used to looked for significant Vanderbilt PRS-KS score differences between the responses of the U.S. representative population and the Vanderbilt graduate-level healthcare students while controlling for age, sex, race, and ethnicity.

## Results

### Development of the Vanderbilt PRS-KS

Developing the *Vanderbilt PRS-KS* was an iterative process of collecting feedback on the scale’s wording, clarity, and comprehensiveness, which involved a team of genetic counselors, geneticists, and medical doctors with considerable clinical genetic experience and education. The team conceived the survey’s content validity with two key domains: benefits of PRSs and limitations of PRSs.

Utilizing existing genetic knowledge scales,^6,22,23^ ten candidate items were piloted for the Vanderbilt PRS-KS (Table S1). The Vanderbilt PRS-KS was designed to represent one latent construct, PRS knowledge. Items addressed misconceptions between PRSs and monogenic tests, which could affect decision-making (e.g., If you receive a high polygenic risk result, your children will have a high polygenic risk result).

### Item Retention

Our analytic plan was to drop items that did not fit the CFA or IRT model, violated local dependence, or violated DIF. Three items (1, 2, and 7) were removed from the ten draft items of the Vanderbilt PRS-KS. We removed draft items one and two based on low standardized factor loadings (Λ < 0.3) in the initial CFA. After rerunning CFA with the remaining eight items, we discovered draft items 7 and 3 displayed local dependence (modification indices > 10). The wording of item 3 (polygenic risk scores are based on genetic changes in more than one gene) and item 7 (a polygenic risk score can be measured at birth) did not reveal surface-level dependence. To promote the IRT assumption of local independence, we individually removed each item and examined the CFA model fit.

Removing item 3 and retaining item 7 led to poor CFA model fit (TLI = 0.945) and introduced new local dependence between items 5 and 10 (modification indices = 10.1). Removing item 7 and retaining item 3 had a good CFA fit with no areas of local dependence. Therefore, we kept item 3 and removed item 7 for analysis, leaving the 7-item Vanderbilt PRS-KS (Table 1).

**Table 1.**
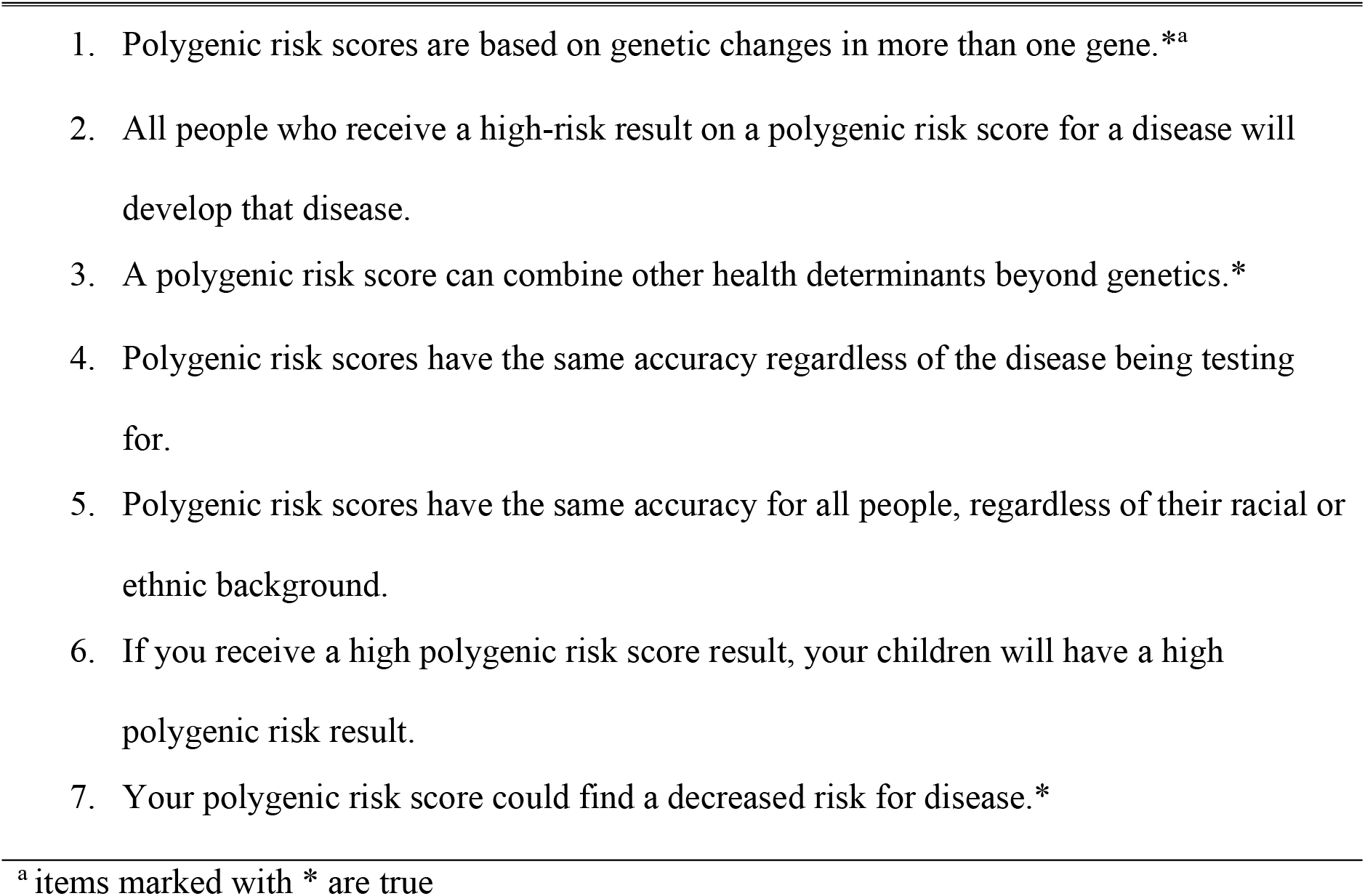
The Vanderbilt PRS-KS.

### Respondents

Five hundred individuals completed the survey using prolific.co’s representative U.S. sample algorithm (Table 2). The percentage of male and female respondents was similar (48.8% and 50.4%), with a majority being white (73.8%) and non-Hispanic (91%). The average age was 45 years old.

**Table 2.** Participant Characteristics.

| Demographics | U.S. Representative Sample<br>(N=500) |  | Graduate-Level Students<br>(N=74) |  |
| --- | --- | --- | --- | --- |
|  | N (%) | Mean (SD) | N (%) | Mean (SD) |
| Age (years) |  | 45 (15.9) |  | 26 (5.38) |
| Sex |  |  |  |  |
| Male | 244 (48.8%) |  | 20 (27%) |  |
| Female | 253 (50.4%) |  | 53 (71.6%) |  |
| Prefer not to answer | 4 (0.8%) |  | 1 (1.4%) |  |
| Race |  |  |  |  |
| White | 369 (73.8%) |  | 60 (81.8%) |  |
| Non-White | 129 (25.8%) |  | 12 (16.2%) |  |
| Prefer not to answer | 2 (0.4%) |  | 2 (2.7%) |  |
| Ethnicity |  |  |  |  |
| Hispanic | 41 (8.2%) |  | 9 (12.2%) |  |
| Non-Hispanic | 457 (91%) |  | 65 (87.8%) |  |
| Prefer not to answer | 4 (0.8%) |  | 0 (0%) |  |

Seventy-four individuals in the VUSN and VUSM completed the survey. The majority were female, 71.6%, white, 81.8%, and non-Hispanic, 87.5%. The average age was 26 years old.

### Descriptive Statistics

The mean proportion of individuals answering each item correctly was 0.59, with a standard deviation of 0.11 (Figure S1). The proportion of correct responses ranged from 0.49 to 0.78, demonstrating no ceiling or floor effects. The item-total correlations were positive and moderate to large magnitude r = 0.35 ranging from 0.16 to 0.55. Similarly, the item-total correlations were positive r = 0.37 with a range of 0.26 to 0.46. The mean summed score on the Vanderbilt PRS-KS was 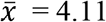 with no identified outliers (Figure S2).

### Confirmatory Factor Analysis

The 7-item Vanderbilt PRS-KS displayed acceptable one-factor confirmatory factor analysis fit (χ^2^ = 22.218, df = 14, p = 0.074, RMSEA = 0.034, CFI = 0.977, TLI = 0.965). The standardized factor loadings showed moderate item contribution to each factor □ = 0.33 to □ = 0.61, suggesting the seven items were associated with the underlying construct-PRS Knowledge (Table S2). There was no evidence of local dependence, and we proceeded with IRT.

### Item Response Theory Analysis

The Vanderbilt PRS-KS fit the 2PL model better than the 1PL model, meaning the items’ location (*b*) and discrimination parameter (α) varied (χ^2^ = 35.14, df = 6, and p < 0.01). We compared the 2PL with the 3PL model to determine if individuals were likely to be guessing. The results indicated that individuals were not likely to be guessing and were selecting don’t know/uncertain as to their response if they were uncertain (χ^2^ < 0.01, df = 1, and p = 1.00). Item-level fit was sufficient for the 2PL model (P > 0.05).

The difficulty parameters ranged from −1.09 to 0.04, indicating modest variation. The discrimination parameters were all positive with a moderate slope (*a* = 0.69 *to a* = 1.95). Therefore, the scale can discriminate between individuals with different levels of PRS knowledge without local dependence.

Item characteristic curves followed an s-shaped curve, increasing as ability increased (Figure S3). Using a false discovery rate of 0.05, there was no evidence of uniform or nonuniform DIF across age, sex, race, and ethnicity subgroups. We proceeded with the 2PL model for IRT scoring.

### Item Response Theory Score and Reliability

For a more straightforward interpretation, the IRT scores were scaled to an estimated T-score metric. Respondents’ IRT T-scores ranged from 29.1 to 65.4 (Figure S4). Table S3 enables practical conversion of summed Vanderbilt PRS-KS scores to expected scaled T-scores. The scale’s TIF demonstrated good reliability (≥ 0.7) for T-Scores between 37.28 and 54.76 (Figure S5). Converted to summed scores, the Vanderbilt PRS-KS showed good reliability (> 0.70) for summed scores ≥ 2 and ≤ 5.

### Classical Test Theory Reliability

The Vanderbilt PRS-KS approaches acceptable reliability in the U.S. representative sample with Cronbach’s α = 0.66. However, the greatest lowest bounds (GLB) demonstrated acceptable reliability (0.70).

The scale’s reliability was poor in the graduate-level healthcare students’ sample (Cronbach’s α = 0.46 and GLB = 0.60).

### Convergent Validity

Pearson’s correlations found evidence of good convergent validity (Table S4). The Vanderbilt PRS-KS moderately correlated with the UNC_GKS genetic knowledge measure (r = 0.55) and slightly correlated with the applied PRS knowledge measure (r = 0.29). The respondents’ attitudes towards genetics did not correlate with the Vanderbilt PRS-KS (p = 0.17).

We tested if the graduate-level healthcare students would perform better on the Vanderbilt-PRS than the U.S. representative population. After controlling for age, sex, race, and ethnicity, multivariate logistic regression found a significant difference in Vanderbilt PRS-KS scores between the graduate-level health care students and the U.S. representative sample (p = 1.6e^-5^). No significant differences were found based on age, sex, race, and ethnicity between the two cohorts.

## Discussion

The present study addresses several limitations from prior studies exploring patients’ PRSs knowledge. First, assessments of PRS knowledge are typically disease-specific, usually cancer.^24–26^ Although cancer PRSs are the most commonly used clinically, PRSs have shown promising clinical utility for various diseases.^27^ We designed the Vanderbilt PRS-KS to be ‘context neutral,’ i.e., not specific to a method of PRS development, disease prediction, or current clinical utility. Therefore, the Vanderbilt PRS-KS’s neutral wording could standardize PRS knowledge across multiple diseases. Second, existing studies assessing individuals’ PRS knowledge did not report the reliability or validity of their knowledge measures.^24,25,28^ Reliability and validity determine how well an instrument measures its intended construct and its consistency across populations and time. Ensuring acceptable reliability and validity of instruments leads to better evidence-based conclusions.

The Vanderbilt PRS-KS was validated using rigorous psychometric methods. IRT has several advantages compared to classical test theory methods, including modeling items individually to the construct measured, allowing for precision estimates across different ability levels of the underlying construct, and providing score estimates independent of the items used.^14^ The Vanderbilt PRS-KS is most sensitive for low to moderate genetic knowledge (summed scores between ≥ 2 and ≤ 5). We expect the general population to fall within one standard deviation of the U.S. representative sample mean, where the Vanderbilt PRS-KS’s results are most precise. Notably, the scale has acceptable reliability for individuals with lower scores (> 0.60), who are more likely to need targeted interventions and support interpreting PRSs.

Another important finding was the Vanderbilt PRS-KS functions similarly, regardless of age, sex, race, or ethnicity. Score differences can be attributed to different levels of PRS knowledge. The Vanderbilt PRS-KS can be administered to diverse demographic groups and produce reliable results.

The present study has several limitations. First, the Vanderbilt PRS-KS’s Cronbach’s alpha value for reliability is in-between acceptable α ≥ 0.7 and poor values α ≤ 0.6. Cronbach’s alpha assumptions of tau-equivalence and normality of the data are often not met and can lead to an under-representation of the true reliability of the scale.^29^ Notably, the scale’s reliability improved to above 0.7 for summed scores between ≥ 2 and ≤ 5. Further, the GLB value of 0.70 was acceptable in the U.S. representative sample population. Second, our factor loading criteria of ≥ 0.3 for item inclusion during CFA was relaxed compared to other suggested methods of ≥ 0.4.^30^ PRS knowledge is a broad construct; therefore, we relaxed the factor loadings to ensure the scale was comprehensive to PRS knowledge. Another suggested factor loading retaining criteria was retaining factor loadings that do not cover zero in their 95% confidence interval, ^30^ which our retained seven items did not cover zero. Finally, for T-scores above 60, the reliability falls below the 0.60 threshold, indicating results for individuals with high PRS knowledge may not be reliable. However, multivariate regression found a significant difference between the Vanderbilt PRS-KS scores in the U.S. representative population and the scores in the graduate-level healthcare students population. Therefore, individuals presumed to have higher background knowledge of PRSs were distinguishable from individuals presumed to possess lower PRS knowledge, even though the scale’s reliability was low. Additionally, individuals with higher PRS knowledge are less likely to need additional PRS education. Thus, the Vanderbilt PRS-KS’s questionable reliability for individuals with higher PRS knowledge should not detract from its utility.

In summary, we provide the Vanderbilt PRS-KS as a promising tool to quantify PRS knowledge in diverse settings. The Vanderbilt PRS-KS is comprehensive while retaining brevity. At seven items, the scale is feasible for clinical and research settings. Adopting the rigorously validated Vanderbilt PRS-KS can standardize the quantification of PRS knowledge and guide future interventions in PRS education.

## Data Availability

The data that support the findings of this study are available on request from the corresponding author, DS.

## Author Contributions

Conceptualization: D.S., A.B., G.H., L.R; Data curation: D.S., Y.L.; Formal analysis: Y.L.; Funding acquisition: A.B.; Investigation: D.S.; Methodology: D.S., A.B., G.H., L.R., Y.L.; Project administration: A.B., G.H.; Software: Y.L.; Supervision: A.B.; Visualization: D.S., Y.L.; Writing-original draft: D.S.; Writing-review & editing: A.B., G.H., L.R.

## Acknowledgements

The study was supported through the Vanderbilt University Medical Center Brock Family Discovery Scholar funding (A.B.) and the Vanderbilt University’s Master of Genetic Counseling Program (D.S.). L.R. is supported by NHGRI grant U01-HG011181. We thank the Vanderbilt University genetic counseling students, medical students, and nursing students who participated in the survey validation.

## Ethics Declaration

The Vanderbilt University Medical Center Institutional Review Board approved this study.

## Notes

### Competing Interest Statement

The authors have declared no competing interest.

