## Supplemental Tables and Figures for "Development and Validation of the *Vanderbilt PRS-KS*, an Instrument to Quantify Polygenic Risk Score Knowledge"

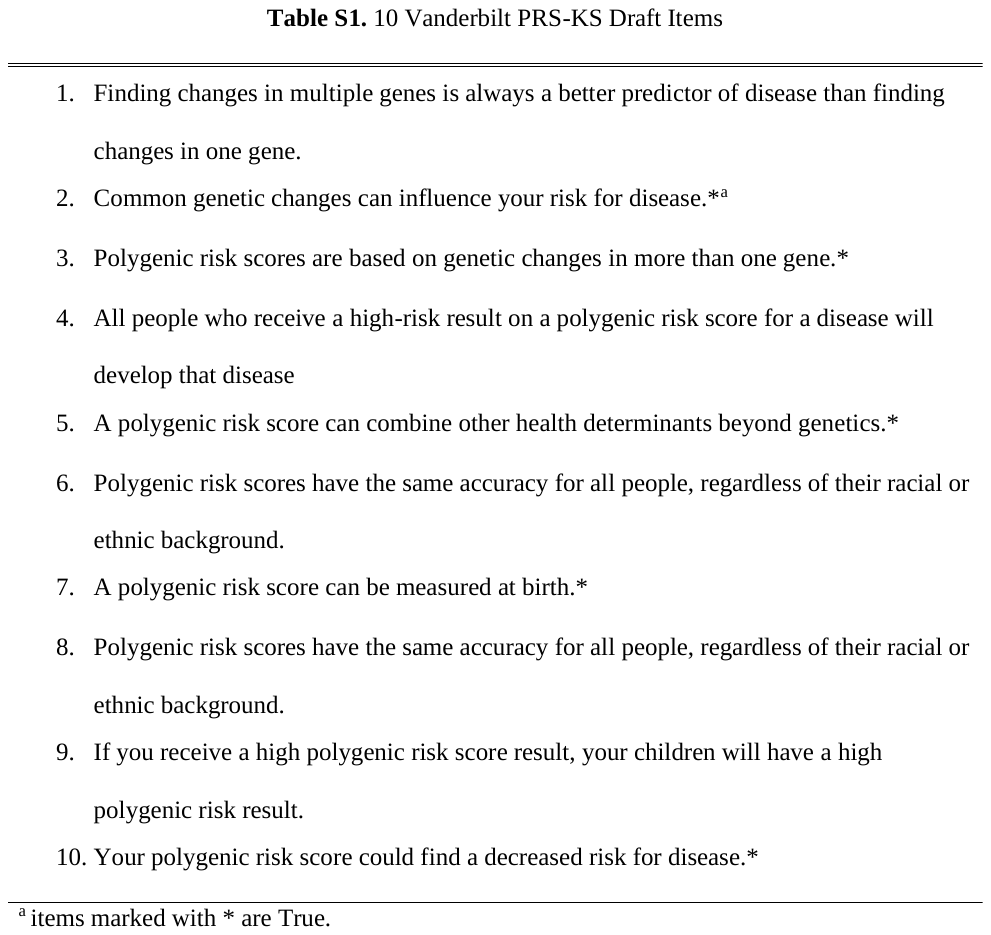

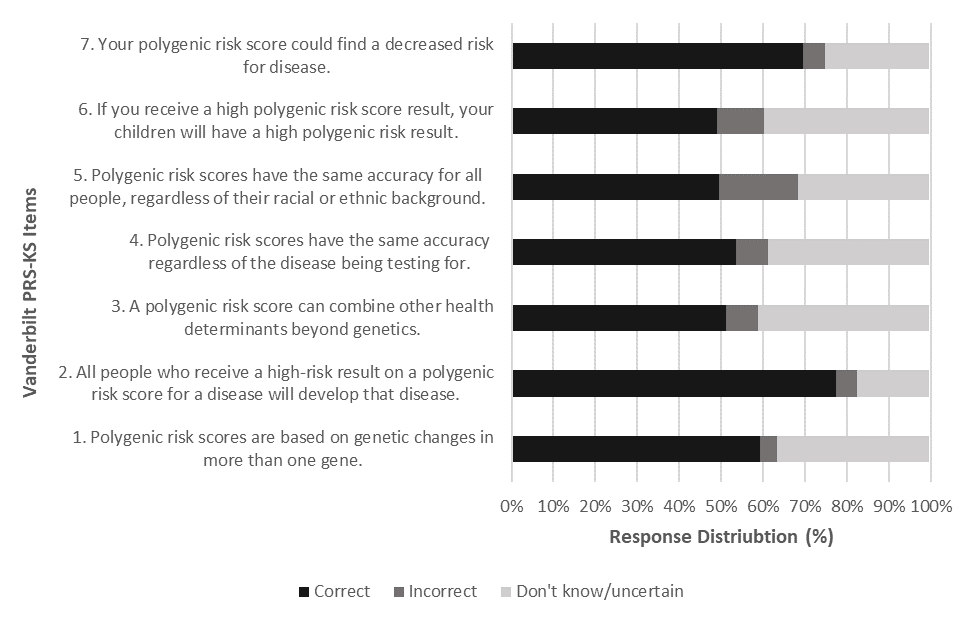
**Figure S1.** Item Response Distribution of the Vanderbilt PRS-KS in a U.S. Representative Sample. There was a total of n = 500 responses. The mean proportion of individuals correctly answering each question was 0.59 (SD = 0.11).

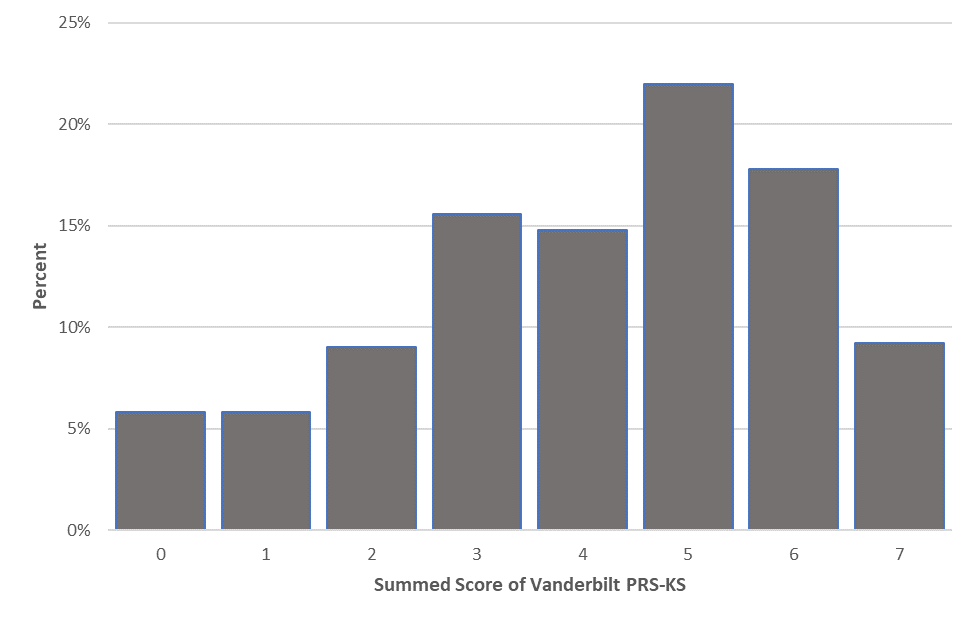
**Figure S2.** Summed Score Distributions of the Vanderbilt PRS-KS in the n = 500 U.S representative sample. The mean score on the Vanderbilt PRS-KS was $\bar{x}$ = 4.11.

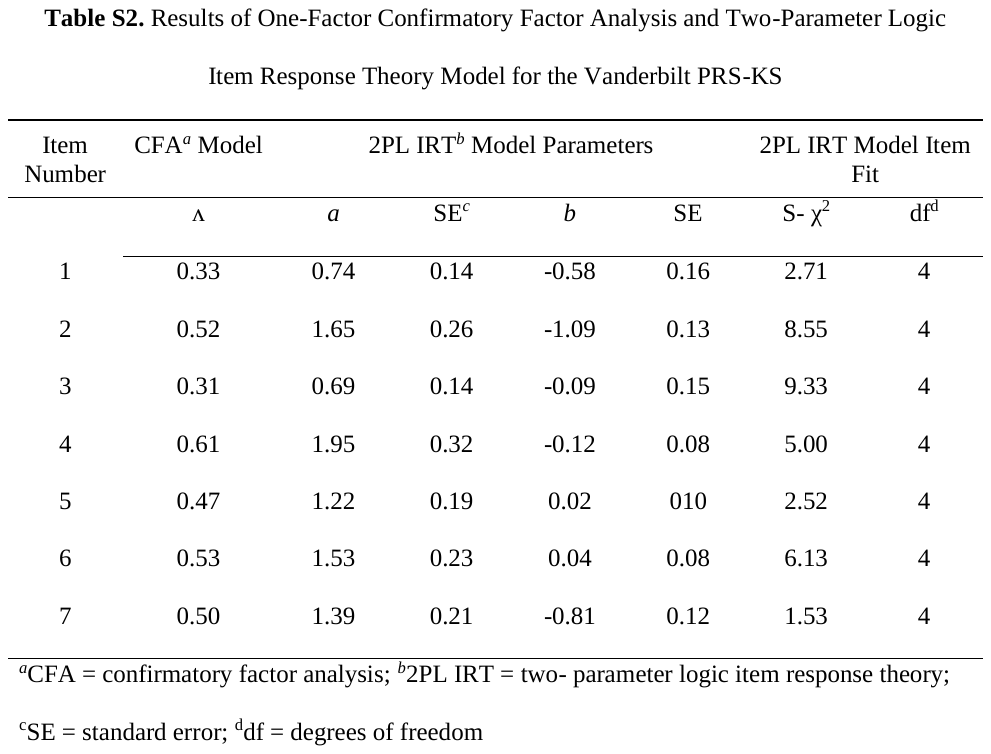

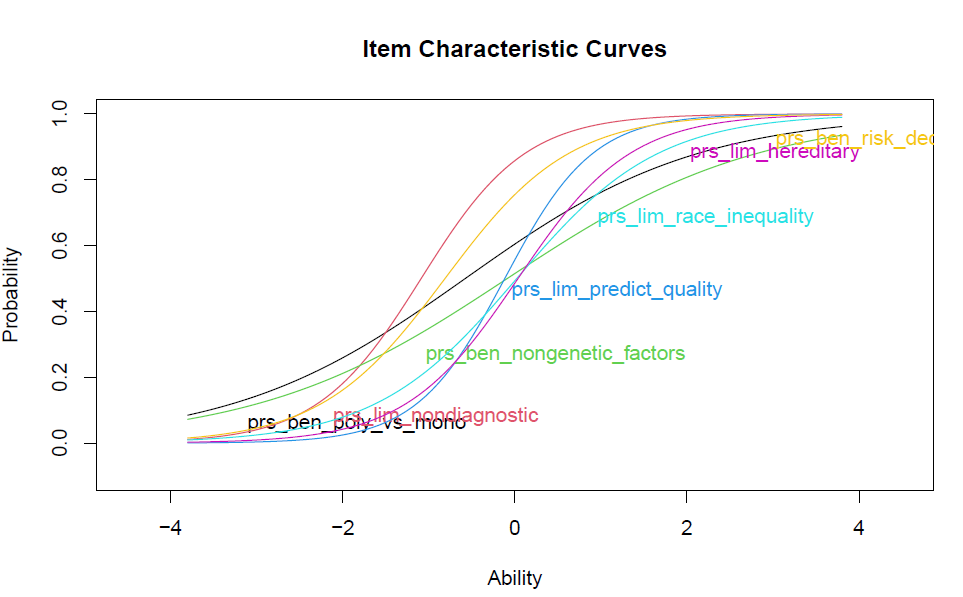
**Figure S3.** Item Response Theory Item Characteristics Curve. Each item on the Vanderbilt PRS-KS has an item characteristic curve labeled with its variable name. The x-axis is the ability level (Θ). Lower ability levels represent lower PRS knowledge levels, and higher ability levels represent higher PRS knowledge. The y-axis indicates the probability of individuals endorsing a question correctly from 0.0 to 1.0. Items shifted to the left on the x-axis are considered easier items (requiring a lower level of PRS knowledge to endorse correctly). In comparison, items to the right are considered more challenging items.

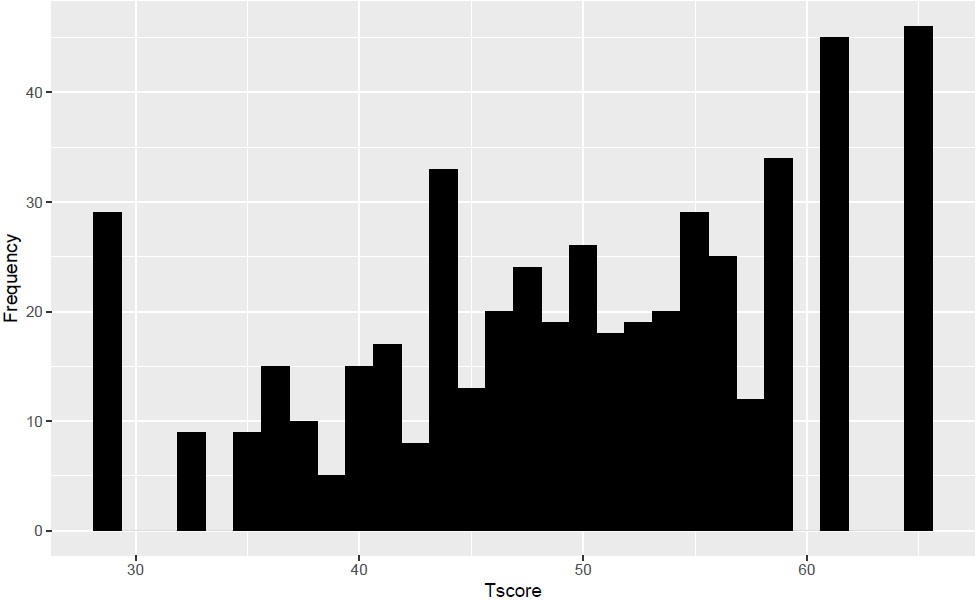
**Figure S4.** Item Response Theory Scores Distribution Scaled to T-Scores. The x-axis represents the summed IRT scores scaled to T-scores, with a population mean of 50 and a standard deviation of 10. The y-axis represents the frequency of the scaled T-score.

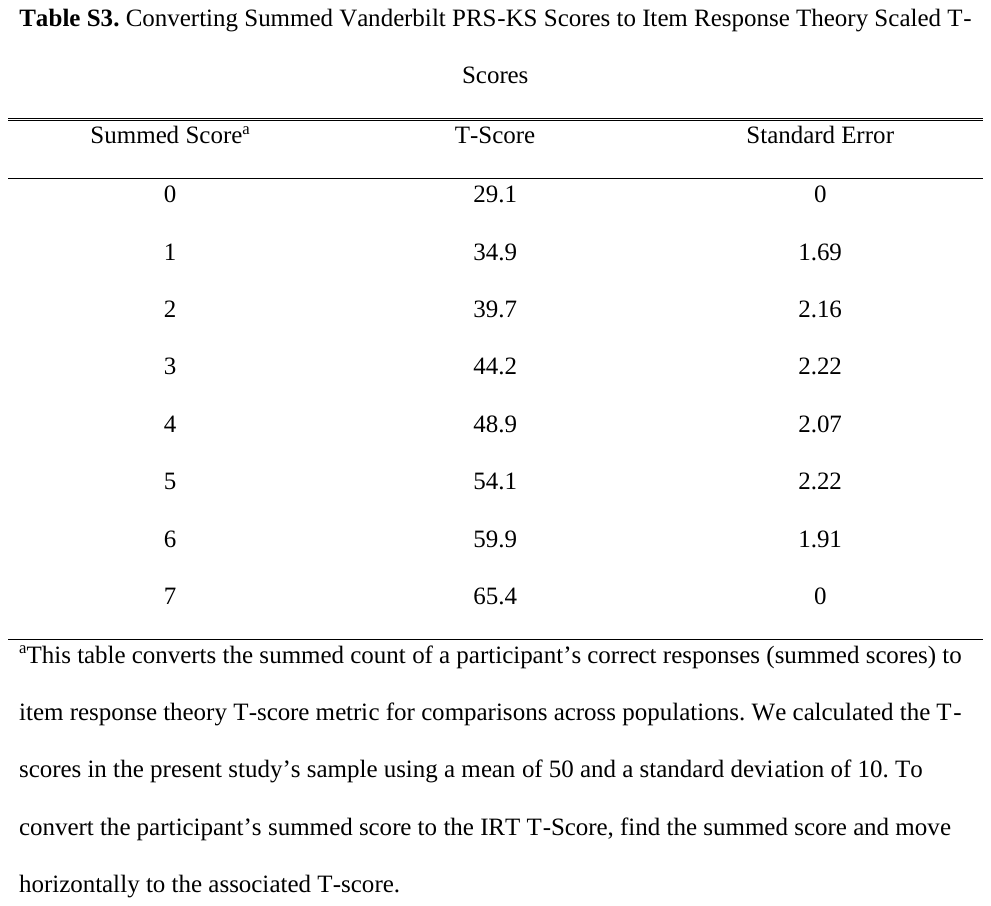

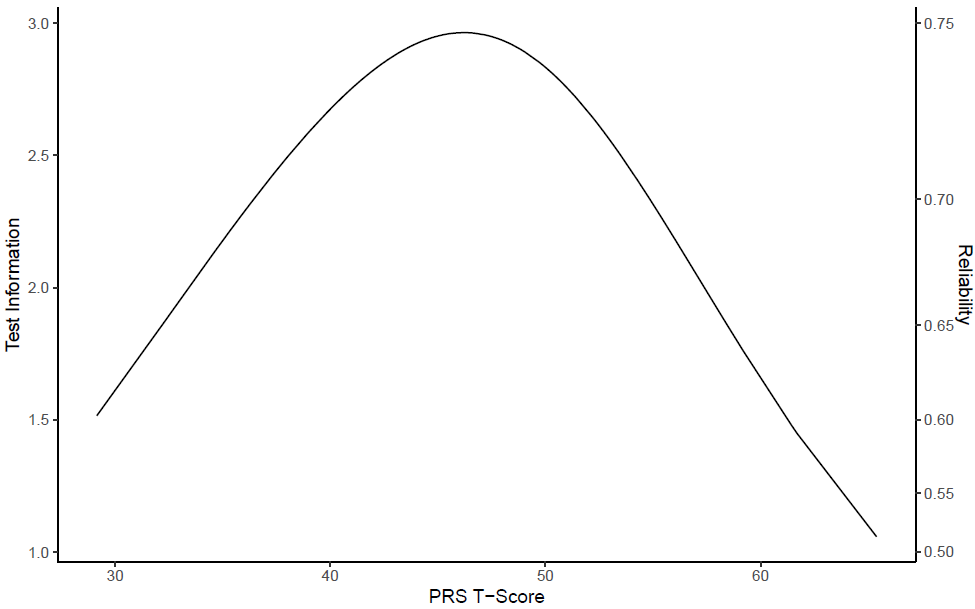
**Figure S5.** Item Response Theory Test Information Curve. The test information curve represents the precision across different levels of PRS Knowledge. The y- axis represents test information, which is the sum of all the items’ information. The x-axis is scaled to represent PRS knowledge through T-scores. The T-scores are scaled to a mean of 50 and a standard deviation of 10.

| **Table S4.** Pearson Correlations Between Vanderbilt PRS-KS and Other Measures | | |
| --- | --- | --- |
| Scale | Correlation With Vanderbilt PRS-KS | 95% CI |
| Genetic Knowledge (UNC_GKS) | 0.55*^a^ | 0.48-0.60 |
| Attitudes Towards Genetics | 0.06 | -0.03-0.15 |
| Applied PRS Measure | 0.29* | 0.21-0.37 |
| ^a^ Items marked with * are P < 0.001; Note: CI = Confidence Interval | | |
